# Implications of genetic heterogeneity for plant translocation during ecological restoration

**DOI:** 10.1101/2020.01.28.923524

**Authors:** Taylor M. Crow, C. Alex Buerkle, Daniel E. Runcie, Kristina M. Hufford

**Affiliations:** Department of Plant Sciences, University of California, Davis, CA, USA; Department of Botany, University of Wyoming, Laramie, WY, 82071, USA; Ecosystem Science and Management, University of Wyoming, Laramie, WY 82071, USA

**Keywords:** *Cercocarpus montanus*, ecological restoration, genetic differentiation, genetic structure, niche, phylogeography, seed transfer, seed transfer zones

## Abstract

Ecological restoration often requires translocating plant material from distant sites. Yet published guidelines for seed transfer are available for very few species. Accurately predicting how plants will perform when transferred requires multi-year and multi-environment field trials and comprehensive follow-up work. In this study, we analyzed the genetic structure of an important shrub used in ecological restorations in the Southern Rocky Mountains called alder-leaf mountain mahogany (*Cercocarpus montanus*). We sequenced DNA from 1440 plants in 48 populations across a broad geographic range. We found that genetic heterogeneity among populations reflected the complex climate and topography across which the species is distributed. We identified several temperature and precipitation variables that were useful predictors of genetic differentiation and can be used to generate seed transfer recommendations. These results will be valuable for defining management and restoration practices for mountain mahogany and other widespread montane plant species.

## Introduction

The restoration of vegetation after a disturbance event can improve many ecosystem services (Barral et al., 2015). For example, soil stabilization, pollinator and wildlife habitat, nutrient cycling, and carbon sequestration are all positively correlated with successful ecological restoration (Benayas et al., 2009). However, bringing foreign plant material to a restoration site can have unintended consequences. Importing maladaptated individuals can result in large-scale plant mortality (Johnson et al., 2004), inbreeding depression of introduced material, or outbreeding depression within future local and foreign hybrid populations (Hufford and Mazer, 2003). Therefore, optimizing the fitness of imported plant material is of vital importance.

Seed transfer guidelines are intended to establish criteria to aid in the selection of plant material for restoration. However, traditional common garden experiments are expensive and time-consuming (Johnson et al., 2004), requiring multi-year and multi-environment field trials and comprehensive follow-up census work. Typically, the relationship between phenotypic variation and environmental and spatial distance are used to create categorical seed transfer zones (Campbell and Sorensen, 1978; Bower and Aitken, 2008), continuous seed transfer guidelines Parker and Niejenhuis (1996), or both (Hamann et al., 2000; Saenz-Romero and Tapia-Olivares, 2008). These experiments however are limited by the number of populations, number of environments, and the amount of time it may take to quantify consequences of importing foreign plant material (Johnson et al., 2004).

Models based on climate (Bower et al., 2014; Crow et al., 2018) or genetic data (Krauss and He, 2006), or a combination of both (Massatti et al., 2020) may be useful for establishing plant population translocation recommendations without the financial or time investment required by a transplant experiment. For example, genetic structure analyses can estimate genetic connectivity between adjacent populations, which can be used to predict the likelihood of reduced fitness during a potential transfer (Ellstrand and Elam, 1993; Sexton et al., 2014). Preserving genetic structure in restoration is important for maintaining adapted combinations of alleles in native populations. Further, gene flow between introduced and native populations may lead to outbreeding depression when locally adapted gene complexes are disrupted by immigrant alleles after admixture (Fenster and Galloway, 2000; Montalvo and Ellstrand, 2001). Identifying geographic and environmental patterns related to genetic differentiation can therefore provide useful guidelines for seed introductions in ecological restoration (Montalvo and Ellstrand, 2001).

In addition to field experiments, knowledge of genetic subdivision of species across space and its association with dimensions of the environmental niche can also contribute to the development of seed transfer guidelines. The niche concept incorporates both the environmental and spatial distribution of a species, and can be used in understanding factors governing range limits (Sexton et al., 2009). One conceptualization of a niche is summarized by Hutchison’s n-dimensional hypervolume (Hutchison, 1957), described as a set of biologically relevant and independent environmental axes within which a species occurs. The multi-variate environmental space represents conditions that accommodate population persistence and growth (Hutchinson, 1978). As habitat quality or availability decreases, population size and gene flow are expected to decrease (Brown, 1984; Eckert et al., 2008). Understanding the relationship between species’ genetic structure and niche can lead to the identification of evolved population differences and locally adapted ecotypes to inform guidelines for seed transfer.

In this study we investigated genetic variation relevant for restoration of a native perennial shrub, alder-leaf mountain mahogany, *Cercocarpus montanus*. Mountain mahogany is used in restoration projects because of its value as a forage plant for large ungulates, especially in the winter months. We collected and sequenced DNA from 1440 individual plant samples from 48 populations, estimated genetic diversity within populations, and measured variation at over 6,000 single nucleotide polymorphisms (SNPs) to describe genetic structure. We tested to what extent genetic structure was a function of latitude, habitat quality, niche centrality, or a combination thereof, with the goal of informing seed transfer recommendations for mountain mahogany.

## Methods

### Study species

*Cercocarpus montanus* Raf. is a deciduous, perennial shrub species in the rose family (Rosaceae) with a large spatial distribution in western North America (Dorn, 2001). The species occurs on both sides of the Continental Divide and from northern Mexico to the Wyoming-Montana state borders in the United States (Fig. 1). Populations are generally distributed between 1200 and 3000 meters in elevation and often grow in rocky, limestone soils (Williams et al., 2004). Mountain mahogany are monoecious and have wind-pollinated flowers. Fruits are achenes with an elongated style that twists in later development, and is covered in trichomes. These structures are hypothesised to aid in wind and animal mediated dispersal (Gucker, 2006). Mountain mahogany shrubs serve as hosts for nitrogen-fixing actinomycete bacteria (genus *Frankia*) in root nodules, and this adaptation contributes to successional processes in arid regions dominated by unstable, low nitrogen soils (Klemmedson, 1979).

**Figure 1:**
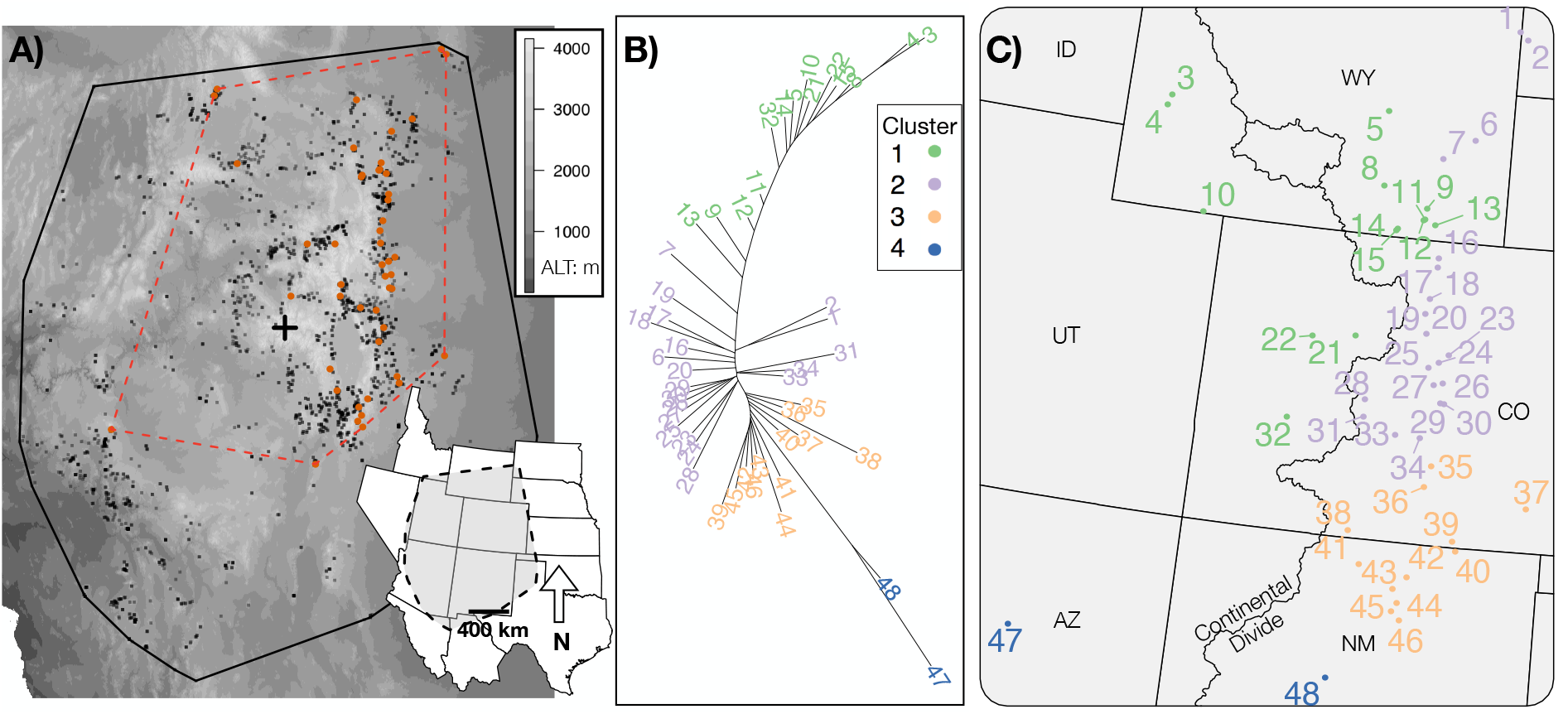
A) Minimum convex polygon (mcp) of species range (black line) around species occurrence points (black squares) and the dashed red line is a mcp around 48 sampled populations (red diamonds). The geographic center of the overall species distribution mcp is marked with a cross. B) Unrooted neighbor-joining tree of Nei’s *D*_*A*_, colors correspond to assigned genetic cluster. C) Map of sampled populations with numbers from 1 to 48 based on latitude for reference.

### DNA extraction, sequencing, assembly and variant detection

Mountain mahogany populations were located along a north-south axis in the southern Rocky Mountains (Fig. 1). We collected leaf tissue from 30 individuals in each of 48 populations and extracted DNA using a modified cetyl-trimethyl ammonium bromide (CTAB) protocol (Doyle, 1987). DNA was quantified with a NanoDrop 2000 spectrophotometer (Thermo Fisher, Inc.), and additional extractions were conducted when necessary due to high levels of contaminants or low DNA concentrations. We prepared genomic libraries for genotype-by-sequencing (GBS) following protocols in Parchman et al. (2012). To summarize, we digested sample DNA with two restriction enzymes (MseI and EcoRI) and ligated barcodes containing unique 8–10 bp sequences to the resulting DNA fragments for each sample to ensure that sequence reads could be assigned to individuals. We then PCR amplified the barcoded restriction-ligation products with standard Illumina primers (1, 5′ - AATGAT-ACGGCGACCACCGAGATCTACACTCTTTCCCTACACGACGCTCTTCCGATCT - 3′; 2, 5′ - CAAGCAGAAGACGGCATACGAGCTCTTCCGATCT - 3′) (Illumina, Inc.).

Barcoded PCR products were combined into two multiplexed libraries of 720 individual samples (with individuals allocated to the libraries randomly to avoid confounding library effects) and sequenced at the University of Texas Genomic Sequencing and Analysis Facility (Austin, Texas, USA) on the Illumina HiSeq 2500 platform using single-end 100 bp reads. After filtering reads for oligonucleotides used in library synthesis and the PhiX genome, with subsequent demultiplexing and assignment of reads to individuals, we had 24000000 sequence reads for further analysis. We completed a de novo genome assembly with a randomly chosen subset of 2.4 × 10^7^ reads using SEQMAN NGEN software (DNASTAR, Inc.). This step resulted in construction of an artificial, partial reference genome containing 111 967 contigs. We used `bwa` (Burrows-Wheeler Aligner; Li and Durbin 2009) to map reads from each individual to this partial reference genome. Once complete, 15 520 448 total reads (64.6%) assembled to the partial reference genome. Aligned reads were then indexed and sorted using samtools and bcftools (Li et al., 2009). We used the command ‘mpileup -P ILLUMINA -u -g -I -f cemo.fasta sorted.bam | bcftools view -N -c -e -g -v -I -d 0.8 −p -P full -t 0.001 -o variants.vcf’ to calculate genotype likelihoods and filter variant sites. We then retained a single SNP per contig and removed SNPs with an allele frequency less than 0.05.

### Population genetic analyses

We estimated genotypes as the mean of the genotype likelihood distribution and constructed a genetic covariance matrix for all individuals. We ran a principal components analysis (PCA) of the genetic covariance matrix using the *prcomp* function in *R* to summarize genetic variation. We tested for correlations between the individual scores on the first two principal component axes and potential drivers of genetic variation such as latitude, elevation, precipitation and temperature. Additionally, genotype data were used to calculate individual admixture coefficients using the sparse non-negative matrix factorization algorithm (sNMF) implemented in the LEA package (Frichot et al., 2014; Frichot and François, 2015) in R. This algorithm estimates ancestry coefficients in a computationally efficient manner. The sNMF algorithm is similar to the program STRUCTURE (Pritchard et al., 2000; Falush et al., 2003), which estimates ancestry independently for each individual, and does not require *a priori* assumptions about population membership. To determine the best-supported number of genetic clusters (K) within our collections of mountain mahogany, we used a cross-entropy criterion from K=1 to K=10 from the snmf function. This criterion uses a masked genotype testing set to determine the prediction accuracy of the model at each K value.

Point estimates of allele frequencies within each population were calculated from the genotype likelihoods, and allele frequencies were used to calculate the Weir moment estimator of F_ST_ (Weir and Hill, 2002) and Nei’s genetic distance (*D*_*A*_) (Nei et al., 1983; Takezaki and Nei, 1996) as measures of genetic differentiation. F_ST_ was calculated using the CALCULATE.ALL.PAIRWISE.Fst function in the BEDASSLE package in R, and *D*_*A*_ was calculated using a custom R script.

We used Bayesian linear models with Nei’s *D*_*A*_ as the response variable, and pairwise geographic distance, environment distance, and a binary variable representing the Continental Divide as model predictors. Population pairs were assigned 0 if they originated from the same side of the Continental Divide, or assigned 1 if they were collected from opposite sides of the divide. Environmental distances were measured as the population pair difference for each environmental variable centered on the mean and divided by the standard deviation (z-score). Environmental variables included thirty-year normal temperature and precipitation estimates from thin plate spline surfaces (http://forest.moscowfsl.wsu.edu/climate). All predictor variables (Table S1) were standardized prior to modeling so that the magnitude of their estimated coefficients could be compared. We fit the full and reduced models for genetic differentiation in R with the rjags package for MCMC models in JAGS (Plummer, 2003). We ran Markov Chain Monte Carlo (MCMC) simulations for 10 000 iterations with he first 2000 steps discarded as burn-in. We thinned the MCMC chain every 5 steps for a total posterior sample of 1600 for each of 3 chains. The deviance information criterion (DIC) was used to select the model that best accounted for genetic distance, as well as to compare models with and without spatial distance, environmental distance, and topographic barriers as covariates.

### Relative contribution of geographic and environmental distance to genetic differentiation

Geographic and environmental distances could contribute to adaptive differentiation that can affect translocation outcomes of seed sources. We used a model that explicitly differentiates between the effects of environment and topographic barriers to gene flow, relative to spatial distance. This model was developed by Bradburd et al. (2013), is called Bayesian Estimation of Differentiation in Alleles by Spatial Structure and Local Ecology, and is implemented in the R package BEDASSLE. We tested the complete dataset, and used the beta-binomial Markov Chain Monte Carlo model. We ran MCMC simulations for 3×10^6^ iterations, thinned the chain every 20 iterations, and checked the trace plots for convergence and acceptance rates.

### Genetic diversity in central and peripheral habitat

We modeled genetic diversity as a function of spatial and environmental centrality. We estimated genetic diversity for each population using the program ANGSD (Korneliussen et al., 2014). Sequence alignments to the pseudo-reference (sorted BAM files) were used as input to calculate each population’s site allele frequencies from genotype likelihoods. We filtered sites that had a minimum mapping quality of 10 and a minimum q-score of 20. The allele frequency likelihoods were used to calculate the maximum likelihood estimate (MLE) of the site frequency spectrum (SFS) using the EM algorithm. Estimates of nucleotide polymorphisms were calculated as *θ*_*π*_ (Tajima, 1983), a measure of average pairwise differences, and Watterson *θ*_*W*_ (Watterson, 1975), which is based on the number of segregating sites. Theta estimates were calculated using the empirical Bayesian approach with the SFS as priors (following http://popgen.dk/angsd/index.php/Thetas,Tajima,Neutrality_tests).

To model spatial and environmental centrality of our collections in the context of the entire range of *C. montanus*, we used range-wide occurrence points from a previous study of mountain mahogany (Crow et al., 2018). Spatial centrality was calculated as the great circle geographic distance (van Etten, 2018) from each of our sampled populations to the mean latitude and longitude of the species’ range, and the range of each individual genetic cluster separately (Fig. 1). We calculated spatial peripherality as the distance between each population and the shortest linear distance to the edge of the minimum convex polygon of the species’ range. Environmental centrality was calculated as the multidimensional euclidean distance of each population to the species’ environmental centroid, and the centroid of each genetic cluster (Blonder et al., 2014). We also used the probability of occurrence derived from a previously published species distribution model (SDM) of mountain mahogany (Crow et al., 2018) as an indicator of habitat quality, which was used as a predictor of population genetic diversity. In summary, environmental variables were selected for the SDM using a model improvement ratio following (Murphy et al., 2010), and a Random Forests algorithm was used to generate the distribution model. We then used linear models to determine to what extent habitat quality, spatial centrality, or environmental marginality were predictive of genetic diversity using the lm and anova function from the stats packages in R.

### Niche similarity among genetic clusters

Niche overlap statistics were used to test if genetic clusters defined by the sNMF admixture analysis occupied distinct subsets of the overall environmental range. Broennimann et al. (2012) developed methods to get an unbiased estimate of niche overlap using kernel smoother functions applied to densities of occurrence points in environmental space, calibrated on the available environmental space across the study area. We calculated kernel densities for the environment occupied by each genetic cluster, and used D metrics (Schoener, 1970) to determine if there was significant overlap of niche space between genetic groups:

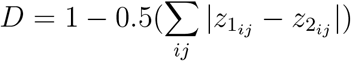

where 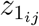 and 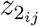 are the occupancy of the environment calculated from kernel density functions of entity one and two respectively. The D metric is 0 if there is no overlap between genetic groups and 1 if there is complete overlap. We used the ecospat package (Broennimann et al., 2017) in R (R Core Team, 2018) to calculate niche similarity and overlap. Ecospat performs a randomization test where *z*1_*ij*_ and *z*2_*ij*_ are combined and randomly separated into two groups, and the D statistic is calculated 100 times to build a null distribution. The observed D statistics, using genetic clusters as entity designations, were calculated and compared to the distribution of simulated D values for each pair of genetic clusters separately. Presence points and environmental data for the distribution of mountain mahogany from Crow et al. (2018) were incorporated as background points.

## Results

### Sequence alignment and SNP discovery

We identified 12 022 single nucleotide variants using samtools and bcftools (Li and Durbin, 2009). For a variant site to be identified, we required that at least 50% of all individuals have a minimum of one read at that locus. After removing sites with a minor allele frequency of <5% and randomly selecting one variant per contig to ensure independence of loci, we retained 6352 single nucleotide polymorphisms (SNPs) for further analyses of population genetic structure. In sum, 1366 of the 1440 individual samples of *C. montanus* had sufficient sequencing coverage to be retained for further analysis, resulting in a range of 22–30 individuals per population. Remaining samples each had an average of 8.5 reads per SNP.

### Population genetic analyses

The first PC axis (PC1) accounted for 89.7% of the genetic variation among individuals of mountain mahogany, and reflected latitude of origin and the effect of the Continental Divide as a barrier (Fig. 2). PC2 accounted for 3.1% of genetic variation, and separated populations of *C. montanus* collected near Albuquerque, NM and Flagstaff, AZ, from those in the remainder of the range. The first PC axis shows that mountain mahogany has continuous genetic variation in the southern portion of its range, and two separate clusters in northern latitudes. Pearson’s correlation coefficients (r) between each environment variable and PC1 were used to determine the likely drivers of population genetic structure. We found that two environmental variables: growing season precipitation (GSP) and the number of degree days less than zero °C (DD0), had the highest correlations (0.44 and 0.42 respectively) with PC1.

**Figure 2:**
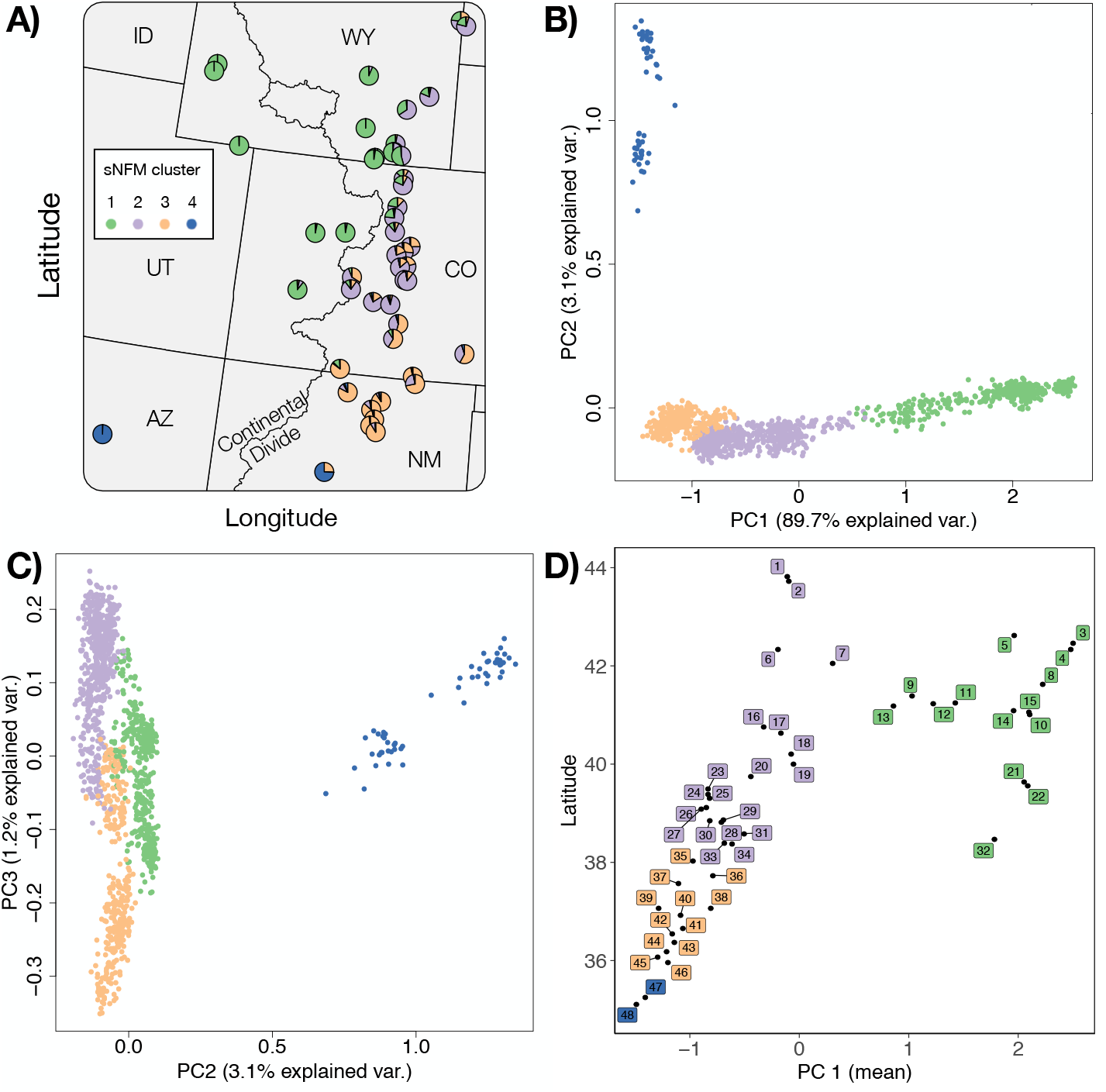
sNFM admixture and principal component analysis of *Cercocarpus montanus*. A) Pie chart of sNFM admixture proportions from k=4 ancestral gene pools (Figure S3) for each of the 48 populations collected in our study. B) PC axis one and two and C) two and three show continuous genetic variation across individuals within clusters. D) Scatter plot of the mean PC axis one score for each of the 48 populations plotted with latitude to visualize geographic structure. Points are colored based on the predominant population assignment from admixture analysis.

The mean Nei’s *D*_*A*_ genetic distance between populations was 0.0346 (SD=0.017), with a range of 0.009–0.108, comparable to previous studies of plant species (Reynolds et al., 2013; Abraham et al., 2015). Pairwise F_ST_ had an overall mean of 0.161, and a SD of 0.0856 (Fig. S1). The mean F_ST_ among pairs of populations from opposite sides of the Continental Divide was 0.241 (SD=0.079), while the mean F_ST_ among populations on the same side of the divide was 0.135 (SD=0.07). Pairwise F_ST_ was positively correlated with spatial distance, and population pairs from opposite sides of the Continental Divide had elevated F_ST_ resulting from the effective topographic barrier (Fig. 3). Growing season precipitation and degree days less than 0°C were standardized and combined as a single mean Euclidean distance for each population pair, and served as environmental predictors in modeling. The Bayesian linear model with the lowest DIC included both spatial and environmental distance as predictors of genetic differentiation (Table 1). The best predictor in a univariate model of genetic differentiation was geographic distance, followed by environmental distance, while the binary design matrix representing the Continental Divide was the worst predictor.

**Figure 3:**
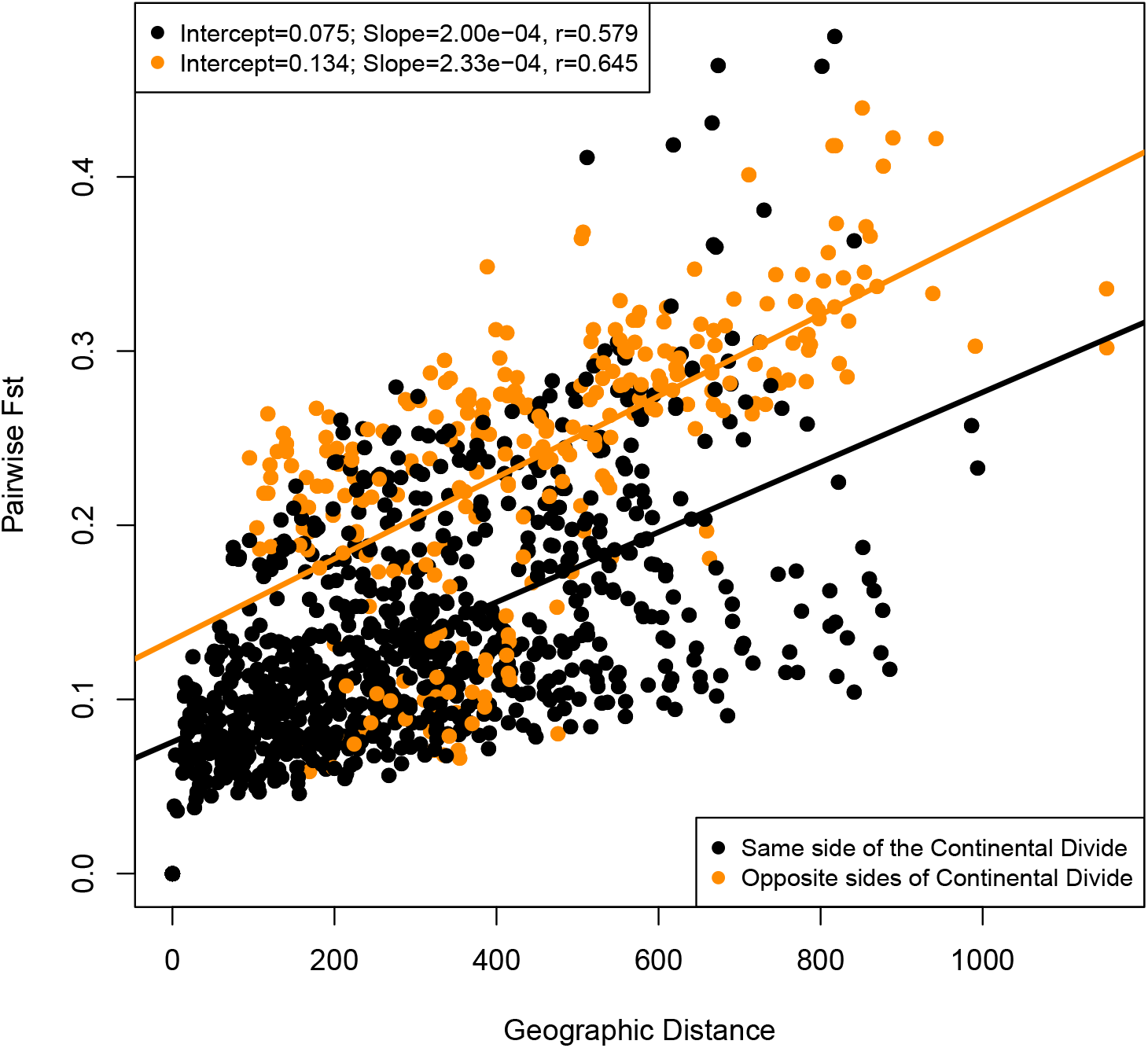
Matrix regression of pairwise genetic and geographic distances. Red points are point pairs from opposite sides of the Continental Divide, while black points are point pairs from the same side of the Continental Divide. Two separate linear models results are listed and model line and summaries correspond to point colors.

**Table 1:** Bayesian linear regression models and coefficients. Predictor variables were standardized using a z-score prior to modeling. Genetic distance (GenDist) was calculated as Nei’s *D*_*A*_. Environmental distance is a multivariate distance matrix of degree days less than zero, and growing season precipitation. Geography is a pairwise geographic distance matrix. The smallest DIC indicates the best model.

| Model | $\beta_1$ (SD) | $\beta_2$ (SD) | $\mu$ (SD) | DIC |
| --- | --- | --- | --- | --- |
| GenDist ~ Environment + Geography | 0.173 (0.013) | 0.297 (0.0126) | -0.0028 (0.0457) | 601.242 |
| GenDist ~ Barrier + Geography | 0.364 (0.0367) | 0.301 (0.0132) | -9.63e-05 (0.0419) | 679.119 |
| GenDist ~ Barrier + Environment | 0.465 (0.0408) | 0.207 (0.0151) | 0.000258 (0.0538) | 921.633 |
| GenDist ~ Geography | 0.327 (0.0135) |  | -0.00132 (0.0494) | 759.724 |
| GenDist ~ Environment | 0.229 (0.0156) |  | 0.00428 (0.061) | 1026.883 |
| GenDist ~ Barrier | 0.533 (0.0435) |  | 0.000125 (0.0491) | 1105.838 |

The best supported number of clusters for sNFM admixture analysis was K=4 (Fig. S2). Populations were assigned to a single cluster based on the predominant population admixture coefficient of individuals within each population (Fig. S3). Although genetic variation is continuous through most of mountain mahogany’s range, the map of admixture composition shows that the genetic clusters are partitioned in geographic space (Fig. 2, panel A), with more highly admixed zones between clusters. The genetic clusters occupied regions of the species environmental space with different multivariate centroids (Fig. 4 panel A). Cluster one and three had no detected overlap in their environmental niche, while clusters one and two and two and three had partial, but not significant overlap in environmental space (Table S3).

**Figure 4:**
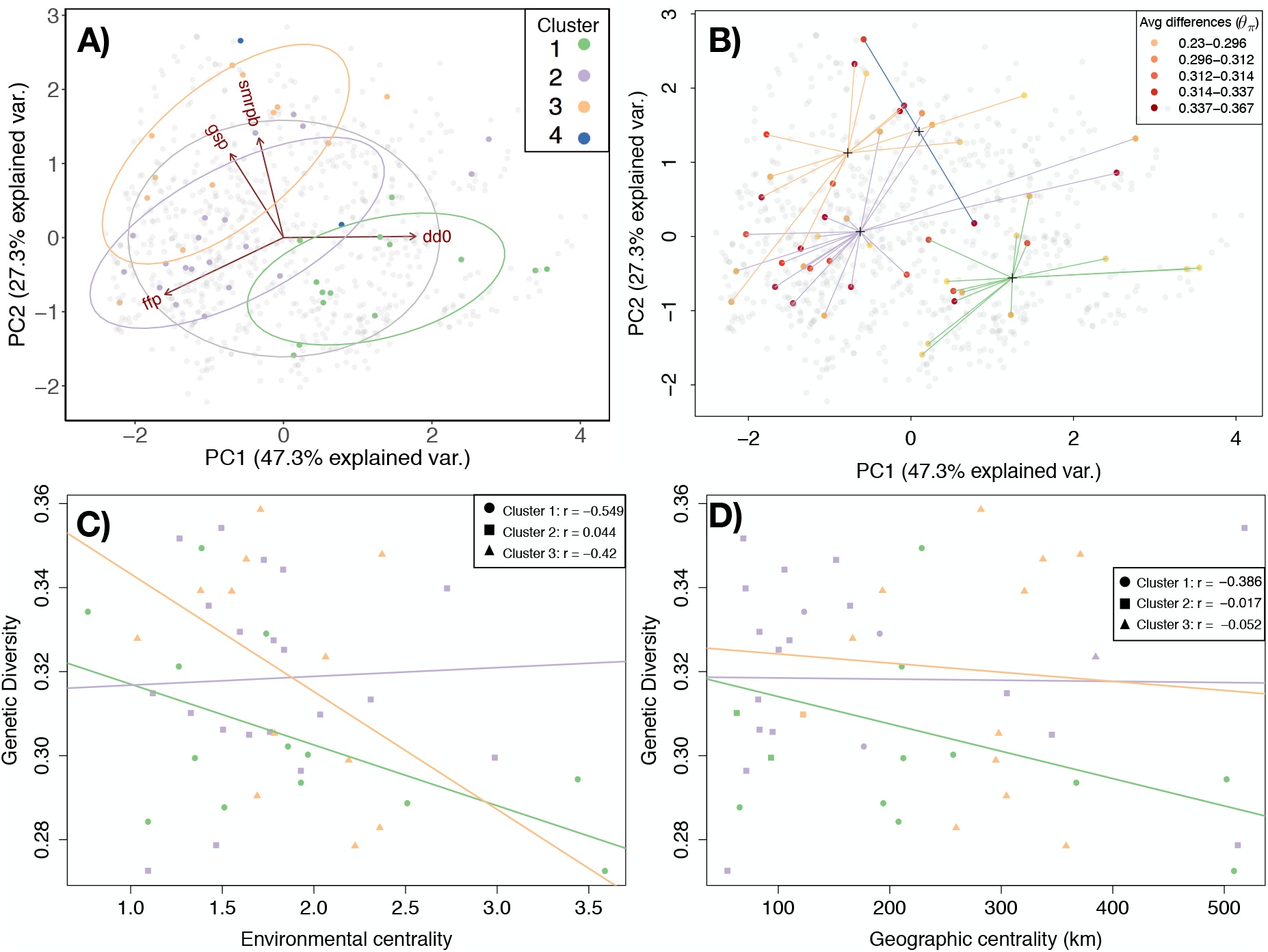
Principal component analysis of growing season precipitation (GSP), summer precipitation balance (smrpb), frost-free period (ffp) and degree days below zero Celsius (dd0). A) The genetic cluster assignment in environmental PCA space, B) shows genetic diversity for each population (point) and the distance (lines) of each population to the cluster-specific environmental centroid (crosses). Finally genetic diversity (*θ*_*π*_) plotted over C) environmental centrality and D) geographic centrality. More central populations are closer to zero. Regression lines were modeled for each genetic cluster separately.

The BEDASSLE analysis calculated the ratio of environmental and spatial distance effect sizes on genetic differentiation (*α*_E_:*α*_D_). We used growing season precipitation and degree days less than 0°C as environmental variables, as well as a binary design matrix representing the Continental Divide to quantify the effect of the environment on genetic distance. A difference of one degree days less than 0°C is comparable to approximately 8 kilometers, and a 1 cm change in growing season precipitation has the same effect on genetic differentiation as approximately 70 kilometers spatial distance. The Continental Divide had the largest effect on genetic differentiation relative to spatial distance. Crossing the Continental Divide had the same effect on genetic differentiation in mountain mahogany as moving 1.7 × 10^7^ km, a larger distance than our collection area.

We detected significant variation in genetic diversity across mountain mahogany’s central range. Nucleotide diversity estimates were highly correlated (r>0.9, *θ*_*π*_ and *θ*_*W*_), and we therefore arbitrarily chose *θ*_*π*_ for further modeling (Table S2). Genetic diversity was not correlated with latitude (P = 0.266, df = 43, *R*^2^ = 0.028). We used two measures of precipitation (growing season precipitation and summer precipitation balance) and two temperature metrics (degree days less than zero and frost-free period) to model genetic diversity because of low colinearity between variables and high correlation with diversity estimates. Genetic diversity was lower in populations farther from the species’ multidimensional environmental centroid. Spatial centrality however, was a poor predictor of *θ*_*π*_. Likewise, we found that spatial centrality was a poor predictor of the probability of occurrence (Fig. S4). The environmental distance to the centroid of each genetic cluster best described genetic diversity, and had a negative correlation (Table 2). We also found significant variation among genetic clusters for the effect of environmental and spatial distance; namely genetic variation within the northern and southern genetic clusters (Cluster 1 and 3) both had a significant relationship to environmental marginality, whereas within the central genetic cluster (cluster 2) diversity was not correlated with environment (Fig. 4).

**Table 2:** Summary of linear regression models and model selection criterion for the effects of geographic and environmental centrality on genetic diversity. *P<0.05,**P<0.01.

| Model | $\beta_1$ (CI) | $\beta_2$ (CI) | $\mu$ (CI) | $R^2$ | adj. $R^2$ | AIC |
| --- | --- | --- | --- | --- | --- | --- |
| $\theta_\pi \sim$ Environment (Env) | -0.01** (-.03 - 0.001) | NA | 0.36 (0.32-0.40) | .106 | .085 | -206 |
| $\theta_\pi \sim$ Geography (Geo) | -0.01 (-0.01 - 0) | NA | 0.31 (0.31-0.32) | 0.033 | 0.010 | -202 |
| $\theta_\pi \sim$ Env + Geo | -.01 (-.03 - 0) | 0 (-0.01 - 0) | 0.36 (0.32-0.40) | 0.109 | .067 | -204 |

## Discussion

Mountain mahogany is increasingly used in restoration programs, particularly because it hosts nitrogen-fixing actinobacteria that allow establishment in nutrient-poor soils, and provides important overwintering forage for wildlife. Despite widespread occurrence in the Rocky Mountain West, no prior ecological genetics study has characterized genetic structure across mountain mahogany’s central range. We sequenced 1440 individuals from six U.S. states in the Southern Rocky Mountains to learn the extent of genetic heterogeneity across the geographic range and the environments occupied by the species.

We found evidence that genetic structure of mountain mahogany was affected by spatial and environmental distance, as well as topographic barriers. The results provide preliminary data for seed sourcing guidelines for mountain mahogany. Genetic variation is important to consider for species management, especially in a restoration setting where hundred or thousands of individual plants are transplanted to a new site (Reed and Frankham, 2003). These results have range-wide implications for mountain mahogany shrubland management, and lay the groundwork for critical decision-making under environmental change.

While genetic structure of mountain mahogany varied continuously across the sampled geographic range, distinct clusters suggest that populations may be adapted to local environmental conditions. However, we cannot infer ecotypic variation in this study based on genetic variation alone. Field studies are needed to determine if individuals have higher fitness within genetic clusters relative to individuals grown at sites outside of their cluster of origin. Despite this limitation, model results suggest that the source of seeds for translocation may affect the viability of the resulting population. The Bayesian model with the best fit included both spatial and environmental distance as factors in population differentiation. Results from the BEDASSLE model, designed to disentangle the effects of spatial and environmental distance, showed that growing season precipitation (GSP) and the number of degree days less than zero (DD0) had large effects on genetic structure in this species. This outcome provides support for seed sourcing guidelines that limit collection to the genetic and the correlated environmental cluster represented by the restoration site.

The Continental Divide is associated with greater genetic differences between Mountain mahogany populations, especially in central Colorado, where the Continental Divide is at its highest altitude. Several studies have shown that the Continental Divide is a strong barrier to gene flow (Schield et al., 2018; Machado et al., 2018). However, to date, no published study has documented this in plant species. Several studies have found significant effects of topographic barriers on genetic differentiation in plant species, including: seas (Jaros et al., 2017), lakes and terrain (Ju et al., 2018), rivers (Geng et al., 2015), mountains (Zhu et al., 2017; Reeves and Richards, 2014), and basins (Bontrager and Angert, 2018). Our data agree with these studies and indicate that populations from opposite sides of the Continental Divide are genetically more isolated, despite what may appear to be close spatial proximity (Fig. 3). Populations from the western slopes of the Rocky Mountains had high among-population genetic differentiation, especially populations 3 and 4 (Fig. 1 panel B and C). Population 3 and 4 may have been founded separately from other western slope populations, or may contain hybrids with a closely related species, *Cercocarpus ledifolius*, that co-occurs in this region (Stutz, 1988). The two most genetically differentiated populations (47 and 48), in New Mexico and Arizona respectively (Fig. 1 panel B and C), inhabit isolated locations surrounded by desert regions with low habitat suitability (Crow et al., 2018). Populations 47 and 48 are likely adapted to high temperature and low precipitation conditions, and may warrant further investigation into their taxonomic status.

Despite the heterogeneity of climatic conditions in our study area, we found that the best supported genetic clusters corresponded to plants in cohesive geographic regions (Fig. 2). Further, the genetic clusters were associated with significantly different environmental space (Fig. 4A), which corroborates linear modeling results showing that spatial distance and environment are both factors related to genetic variation. Given these results, we analyzed patterns of genetic diversity across both spatial and environmental gradients.

Model outcomes suggested that environmental centrality was a better predictor of genetic diversity than spatial distance. This analysis was completed for all sampled populations, as well as for individual genetic clusters. In both cases, genetic diversity was lower near the environmental niche periphery and not strongly correlated with geographic centrality. A previous study by Lee-Yaw et al. 2017 found similar results, where genetic diversity of *Ara-bidopsis lyrata* ssp. *lyrata* was lower at the edge of the environmental niche, but not the limits of the sampled geographic range. Several meta-analyses have shown that the geographic and environmental range limits do not necessarily coincide, and that the geographic range frequently does not explain patterns of genetic variation (Eckert et al., 2008; Pironon et al., 2017). Another review by Lira-Noriega and Manthey (2014) found that only about half of species ranges have any correlation between geographic and environmental marginality, and that environmental marginality was consistently associated with genetic diversity, while geographic marginality was not.

Reduced genetic variation associated with range limits does not distinguish whether populations occurring at range limits are demographic sinks maintained by immigration from more central habitat, or are important genetic resources adapted to marginal conditions by selection. However, the correlation of genetic diversity and environmental centrality bolsters our findings of genetic structure covarying with the environment. The lack of genetic homogeneity in mountain mahogany indicates that populations are not equivalent, and caution 353 should be taken when planning transfer of plant propagules, particularly during restoration. Other studies of genetic variation near range limits have found contrasting results, even among populations of a species. For example, Hargreaves and Eckert (2018) found that while some populations of the annual plant *Rhinanthus minor* near the range margin had lower fitness, other edge populations were locally adapted. Aguirre-Liguori et al. (2017) found that genetic diversity was lower near the geographic range margin of teosinte, and candidate adaptive SNPs were positively correlated with distance to niche centroid, arguing that populations near the geographic range margins were isolated, while populations near the edges of the environmental niche were locally adapted. In *Picea sitchensis*, populations proximal to the range margin are more likely to carry rare alleles (Gapare et al., 2005), however, a second study of *P. sitchensis* determined that populations near the range limit were locally adapted (Mimura and Aitken, 2010). These studies illustrate that range margins can harbor both source and sink genetic pools even within species, and that making predictions about population fecundity near range margins is difficult.

The results of our study suggest that populations of mountain mahogany have genetic structure across its range that is correlated with differences in the environment. The effect of the Continental Divide on genetic structure was significant. This suggests that transferring populations across the Continental Divide would increase the likelihood of maladaptation, and subsequent risks for outbreeding depression among progeny of local and introduced plants. Two climate variables, degree days less than zero and growing season precipitation, were significantly related to population genetic structure as well as differences in genetic diversity. These two variables could delimit collection sites when transferring seed sources during restoration. Choosing a commercial seed source or collection location that is most environmentally similar to the restoration site may increase chances of introducing adapted genotypes (Hufford and Mazer, 2003). In the case of mountain mahogany, preliminary seed collection zones could be delineated by the four common clusters in genetic analysis. This is a practical approach given that the four clusters represent large spatial regions for collection despite considerable altitudinal and microhabitat variation. Whether populations near range margins are important resources for conservation in mountain mahogany remains unclear. Plants are subjected to biotic and abiotic stressors that influence population dynamics (Pagel and Schurr, 2012; Franklin et al., 2016), seed predators (Louda, 1982), pollinators (Biesmeijer et al., 2006), and dispersers (Merow et al., 2011). As a result, additional studies are needed to determine the adaptive value of mountain mahogany populations along range margins for ecological restoration, particularly in light of changing climate conditions.

## Acknowledgments

We are grateful for the assistance of personnel at New Mexico State University, the USDA Manitou Experimental Forest, Boulder County Parks and Open Space and Mountain Cement Co. in Laramie, Wyoming. This material is based upon work that is supported by Wyoming Agricultural Experiment Station funding provided through the National Institute of Food and Agriculture, U.S. Department of Agriculture, McIntire-Stennis under #228001. Work was also supported by The Berry Biodiversity Center research grant, and Boulder County Parks and Open Space research grant. We would like to also thank P. Mcllvenna and D. Bergman for help collecting leaf material, and L. Mandeville for assistance with DNA library preparation.

