## Supplemental for "Implications of genetic heterogeneity for plant translocation during ecological restoration"

### 1 Tables

#### 1.1 Table S1

**Table 1:** Complete list of climate rasters downloaded from Moscow Forestry Sciences Laboratory.

| Raster layer | Description |
|---|---|
| D100 | Julian date the sum of degree-days >5 degrees C reaches 100 |
| DD0 | Degree-days <0 degrees C (based on mean monthly temperature) |
| DD5 | Degree-days >5 degrees C (based on mean monthly temperature) |
| FDAY | Julian date of the first freezing date of autumn |
| FFP | Length of the frost-free period (days) |
| GSDD5 | Degree-days >5 degrees C accumulating within the frost-free period |
| GSP | Growing season precipitation, April to September |
| MAP | Mean annual precipitation |
| MATTENTHS | Mean annual temperature |
| MMAXTENTHS | Mean maximum temperature in the warmest month |
| MMINTENTHS | Mean minimum temperature in the coldest month |
| MMINDD0 | Degree-days <0 degrees C (based on mean minimum monthly temperature) |
| MTCMTENTHS | Mean temperature in the coldest month |
| MTWMTENTHS | Mean temperature in the warmest month |
| SDAY | Julian date of the last freezing date of spring |
| SMRP | Summer precipitation: (Jul+Aug) |
| SMRPB | Summer precipitation balance: (Jul+Aug+Sep)/(Apr+May+Jun) |
| SMRSPRPB | Summer/Spring precipitation balance: (Jul+Aug)/(Apr+May) |
| SPRP | Spring precipitation: (Apr+May) |
| WINP | Winter precipitation: (Nov+Dec+Jan+Feb) |
| ADI | Annual dryness index, DD5/MAP |
| SMI | Summer dryness index, GSDD5/GSP |
| PRATIO | Ratio of summer precipitation to total precipitation, GSP/MAP |
| AHMI | annual heat:moisture index, ((MAT+15)/MAP)*10 |
| DEM | Digital elevation model |

#### 1.2 Table S2

**Table 2:** Nucleotide diversity and neutrality statistics for populations of mountain mahogany. Populations are numbered by the latitude which they were collected, starting with number one as the most northerly location.

| Population | $\theta_W$ | $\theta_\pi$ |
|---|---|---|
| 1 | 0.354 | 0.354 |
| 2 | 0.218 | 0.279 |
| 3 | 0.224 | 0.273 |
| 4 | 0.281 | 0.294 |
| 5 | 0.248 | 0.294 |
| 6 | 0.257 | 0.305 |
| 7 | 0.271 | 0.315 |
| 8 | 0.299 | 0.3 |
| 9 | 0.39 | 0.349 |
| 10 | 0.189 | 0.23 |
| 11 | 0.3 | 0.299 |
| 12 | 0.349 | 0.321 |
| 13 | 0.24 | 0.284 |
| 14 | 0.271 | 0.289 |
| 15 | 0.339 | 0.329 |
| 16 | 0.348 | 0.336 |
| 17 | 0.356 | 0.347 |
| 18 | 0.353 | 0.344 |
| 19 | 0.256 | 0.306 |
| 20 | 0.298 | 0.31 |
| 21 | 0.288 | 0.288 |
| 22 | 0.356 | 0.334 |
| 23 | 0.309 | 0.329 |
| 24 | 0.368 | 0.352 |
| 25 | 0.201 | 0.273 |
| 26 | 0.299 | 0.313 |
| 27 | 0.258 | 0.296 |
| 28 | 0.319 | 0.34 |
| 29 | 0.274 | 0.306 |
| 30 | 0.31 | 0.325 |
| 31 | 0.232 | 0.3 |
| 32 | 0.299 | 0.302 |
| 33 | 0.329 | 0.327 |
| 34 | 0.268 | 0.31 |
| 35 | 0.324 | 0.328 |
| 36 | 0.378 | 0.339 |
| 37 | 0.269 | 0.299 |
| 38 | 0.221 | 0.283 |
| 39 | 0.387 | 0.359 |
| 40 | 0.301 | 0.305 |
| 41 | 0.246 | 0.29 |
| 42 | 0.359 | 0.339 |
| 43 | 0.389 | 0.347 |
| 44 | 0.207 | 0.278 |
| 45 | 0.361 | 0.348 |
| 46 | 0.313 | 0.323 |
| 47 | 0.261 | 0.328 |
| 48 | 0.349 | 0.367 |

##### 1.3 Table S3

**Table 3:** Niche overlap D statistic. No significant overlap was found between the niche occuppied by any of the three major genomic clusters in this study.

| Zone comparison | D statistic | p-value |
|---|---|---|
| One vs. Two | 0.0798 | 0.545 |
| One vs. Three | 0.301 | 0.336 |
| Two vs. Three | 0 | 1 |

#### 2 Figures

##### 2.1 Figure S1

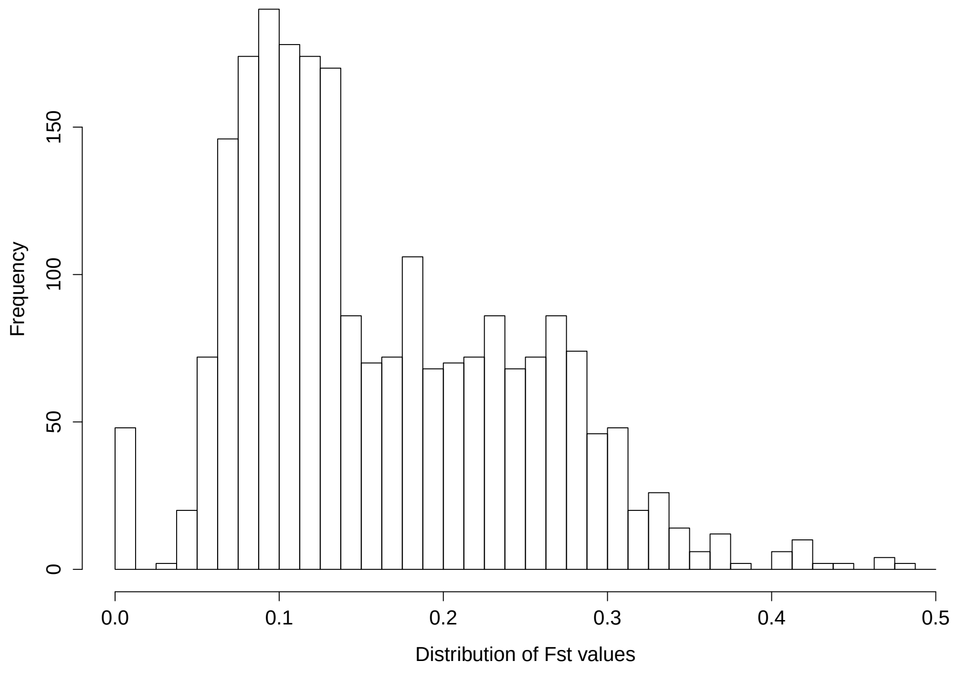

**Figure 1:** Distribution of $F_{ST}$ values among populations of *Cercocarpus montanus* in our study. Notice that the data are skewed, likely due to the effect of isolation by barrier.

#### 2.2 Figure S2

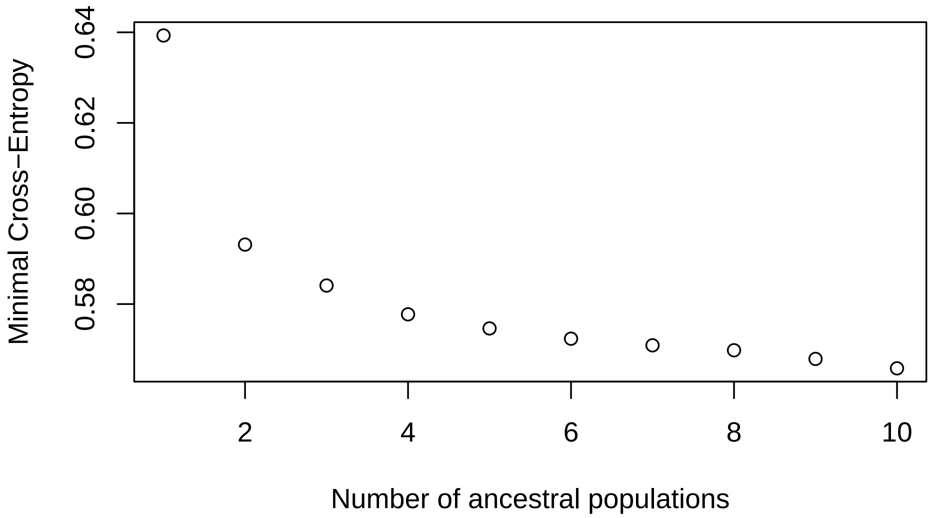

**Figure 2:** The minimum cross-entropy scores for K values 1-10.

#### 2.3 Figure S3

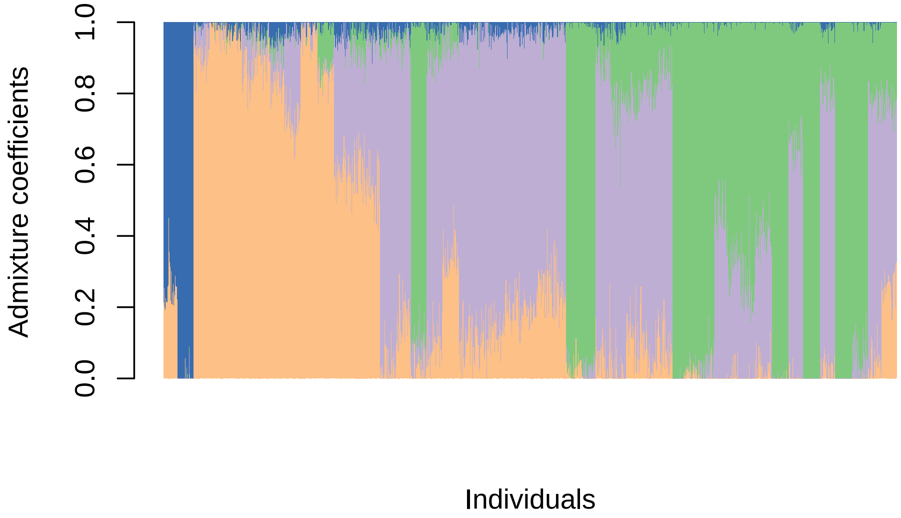

**Figure 3:** Barplot of sNFM admixture results for all 1309 individuals in this study. Samples are arranged by latitude.

#### 2.4 Figure S4

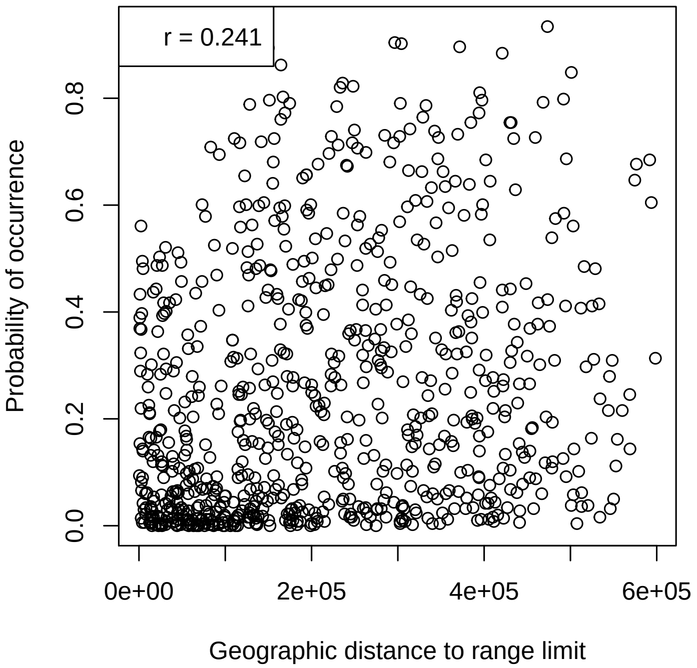

**Figure 4:** Randomly sampled one thousand points across the range of *Cercocarpus montanus* to determine the relationship between the probability of occurrence and the geographic distance to range limit. Very weak positive relationship indicates that in heterogeneous environments, the distance to the range margin does not have a strong relathionship with the quality of habitat.
